# Cross-ecosystem transcriptomics identifies distinct genetic modules for nutrient acquisition in maize

**DOI:** 10.1101/2020.09.02.269407

**Authors:** Yusaku Sugimura, Ai Kawahara, Hayato Maruyama, Tatsuhiro Ezawa

**Author notes:** Corresponding author: Tatsuhiro Ezawa. Author Contributions: H.M., A.K., and T.E. conceived the project; A.K. and T.E. designed the field surveys; Y.S., H.M., A.K., and T.E. collected samples; Y.S., H.M., and A.K. extracted RNA and analyzed plant and soil samples under the supervision of T.E.; Y.S. performed quality assessment and mapping of the sequenced reads; Y.S. and T.E. analyzed the data and interpreted the results; Y.S. and T.E. wrote the manuscript with input from all authors; and all authors discussed the manuscript.

## Abstract

Plants have evolved diverse strategies for the acquisition of the macro-nutrients phosphorus and nitrogen; e.g., mycorrhizal formation, root development, and secretion of chelators/hydrolases to liberate inorganic phosphate. Despite the extensive studies on the individual strategies, there is little information about how plants regulate these strategies in response to fluctuating environment. We approached this issue via profiling transcriptomes of plants grown in large environmental gradients. Roots, leaves, and root-zone soils of 251 maize plants were collected across the US Corn Belt and Japan. RNA was extracted from the roots and sequenced, and the leaves and soils were analyzed. Nineteen genetic modules were defined by weighted gene coexpression network analysis and functionally characterized according to gene ontology analysis, by which we found three modules that are directly involved in nutrient acquisition: mycorrhizal formation, phosphate-starvation response (PSR), and root development. Correlation analysis with soil and plant factors revealed that both phosphorus and nitrogen deficiencies upregulated the mycorrhizal module, whereas the PSR module was upregulated mainly by deficiency in phosphorus relative to nitrogen. Expression levels of the root development module were negatively correlated with those of the mycorrhizal module, suggesting that nutrient acquisition through the two pathways, mycorrhizas and roots, are opposite strategies that are employed under nutrient-deficient and -enriched conditions, respectively. The identification of the soil and plant factors that drive the modules has implications for sustainable agriculture; activation/optimization of the strategies is feasible via manipulating the factors. Overall, our study opens a new window for understanding plant response to complex environments.

## Introduction

Intensification of global food production to meet the requirements of a growing population with minimizing environmental impacts, that is, sustainable intensification of agriculture (1, 2) is a major challenge in our society. Conventional agriculture that heavily relies on chemical fertilizer produced from non-renewable resources, e.g., phosphate rock (3), has contributed to feeding today’s population of the world, but posed threats to ecosystem structure and function, e.g., deterioration of air and water quality (4, 5). In addition, nutrient enrichment also has a negative impact on soil function. Urea application reduces nitrogen (N) mineralization from organic matter through switching N source of microorganisms from organic matter to mineral N (6). High-input of chemical fertilizer generally reduces the abundance and diversity of mycorrhizal fungi (7). Various approaches for reducing chemical fertilizer input have been proposed: the enhancement of nutrient-acquisition efficiency in crops by selection and genetic manipulation (8, 9) and acceleration of nutrient cycling through managing cropping systems (10) and mycorrhizal associations (11).

Plants have developed diverse strategies for the acquisition of the macronutrients N and phosphorus (P) during evolution. N availability has a strong impact on root development. Moderate N deficiency promotes root development in *Arabidopsis* (12). N (nitrate and ammonium)-enriched patches induce localized proliferation of lateral roots for efficient capture of the nutrients (13), which is triggered by nitrate and ammonium uptake via plasma membrane NITRATE TRANSPORTER 1 (14, 15) and AMMONIUM TRANSPORTER 1;3 (16), respectively. Lateral root formation is, however, restricted under severe N deficiency, which may be an important trait for survival in resource-limited environments (17). Localized proliferation of lateral roots was also observed in patches of inorganic phosphate (Pi) (13). Pi deficiency generally inhibits primary root growth, but increases lateral root growth and density, leading to a shallow root system (12, 18). Root hairs also significantly contribute to Pi uptake via extending the surface area for Pi uptake (19). Pi-starvation responses (PSR) have been studied extensively in plants, which include the upregulation of high-affinity Pi transporter genes of the PHT1 family, secretion of non-specific acid phosphatase for mineralization of organic P, excretion of organic acids for the chelation of the counter cations (e.g., Al^3+^, Ca^2+^, and Fe^3+^) of Pi, and replacement of phospholipids with sulfolipids and galactolipids for accelerating internal Pi recycling (20).

Association with arbuscular mycorrhizal (AM) fungi is a widespread strategy in land plants; the fungi facilitate an extensive surface area for the acquisition of mineral nutrients, in particular P and N (21). Deficiencies in P and N trigger secretion of the plant hormone strigolactones into the rhizosphere (22), which facilitates contact of fungal hyphae with the roots via stimulating hyphal branching (23). After the physical contact, the fungal hyphae grow into the cortex and form the highly-branched hyphal termini “arbuscules” where nutrient exchange between the symbionts occurs; briefly, Pi, nitrate, and ammonium are taken up from the soil by extraradical hyphae, delivered to the arbuscules, and released into the arbuscular interface (reviewed in 24) from which the plant cell takes up the nutrients via the mycorrhiza-specific transporters for Pi (25, 26), nitrate (27), and ammonium (28). In return, the host supplies lipid (29-32) and sugar (33) as carbon source for the fungi via the complex of half-size ABC transporters STR/STR2, a putative lipid exporter (34), and the sugar transporter SWEET (35), respectively.

Despite the extensive studies on the individual strategies for nutrient acquisition, there is little information about how plants modulate resource allocation to these strategies in response to biotic and abiotic factors, e.g., nutrient availability, soil physical and biological properties, water availability, and agricultural practice. Optimization of these plant-intrinsic strategies for nutrient acquisition would enhance soil nutrient cycling, which could be one solution for the reduction of fertilizer input. To this end, understanding of the regulatory mechanism of the strategies under field conditions is necessary. We approached this issue via analyzing transcriptomes of plants grown in large environmental gradients, that is, cross-ecosystem transcriptomics. Recently, transcriptomics has been applied to field plants for dissecting their responses to fluctuating environments (36-39). For example, two sorghum genotypes with different drought tolerance showed distinct transcriptomic responses to drought stress, including the rapid upregulation of photosynthetic genes during drought recovery in a tolerant genotype and the downregulation of AM-associated genes during drought stress in both genotypes (36). Even though these studies were conducted only in one field site, their approach opened a new window for understanding plant responses to environmental changes and agricultural practice, which could not be simulated in growth chamber and greenhouse.

Maize (*Zea mays* L.) is one of the most important cereals worldwide, serving as staple food, livestock feed, and industrial raw material (40). We collected maize samples across the US Corn Belt where the world’s highest productivity is achieved by the typical high-input agricultural system, as well as across Japan where agricultural practice is regionally varied due to diversity in climate and soil properties. These large environmental gradients among the sampling sites enabled us to define distinct genetic modules for nutrient acquisition and to identify soil/plant factors that drive the expression of the modules and their interplays.

## Results and Discussion

### Field sites and RNA sequencing

We collected root, leaf, and root-zone soil samples from 66 plants grown in five field sites across the US Corn Belt in 2017 and from 185 plants grown in seven field sites across Japan in 2018, including 17 experimental plots in five out of the seven Japanese sites (Figs. 1A and B, Dataset S1, and SI Appendix, Table S1). The sites and plots were characterized by a principal component (PC) biplot with the climatic/soil factors from which one of the soil/plant factors that were highly correlated with each other (|*r*| ≥ 0.9) were excluded (Fig. 1C and Dataset S2). The US sites are localized in the first quadrant, whereas the Japanese sites are localized in the other quadrants, between which annual precipitation and soil properties were largely different. Among the Japanese sites, the contents of soil organic matter, base, and clay were highly variable. Notably, there were large gradients of N) and P availability among the sites: NO_3_-N, 10.5 – 212 mg kg^-1^; NH_4_-N, 1.0 – 96 mg kg^-1^; Bray II-P, 5.3 – 494 mg P kg^-1^, which led us to the expectation that there would also be large gradients in plant responses to nutrient deficiency/excess.

**Figure 1.**
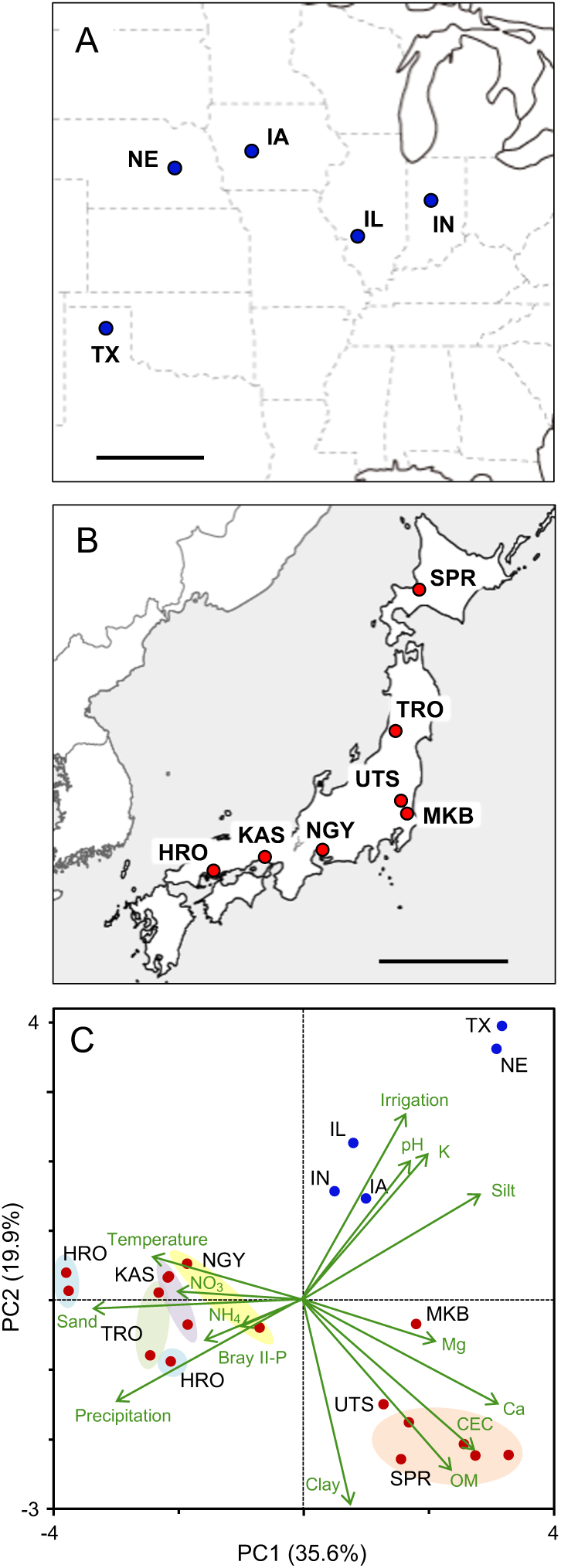
Location and characterization of the maize sampling sites. Sampling sites in the USA (*A*) and Japan (*B*). (*C*) Principal component analysis of soil, plant, climatic, and agricultural variables in 22 plots of the sampling sites in the USA and Japan. Different experimental plots within a site are indicated with colored circles. Detailed information of the plots/sites is described in SI Appendix, Table S1 and Dataset S1. IL, Illinois; IN, Indiana; IW, Iowa; NE, Nebraska; TX, Texas; SPR, Sapporo; TRO, Tsuruoka; UTS, Utsunomiya; MKB, Makabe; NGY, Nagoya; KAS, Kasai; HRO, Hiroshima.

We extracted RNA from all root samples, performed 75-base single end sequencing on a Illumina platform, and mapped the sequence reads to the maize genome (41), by which 8.4 ± 0.25 million reads per sample in average were mapped to exons (Dataset S3). The raw read counts were normalized to Transcripts Per kilobase Million (TPM), and only genes of which the average TPM was 5 or more were subjected to subsequent analysis.

### Definition of gene-coexpression modules

In preliminary analysis, we focused on the genes involved in AM formation because a number of plant genes that facilitate AM fungal colonization, in particular those involved in arbuscule development/functioning, are highly conserved in the plant kingdom (42), implying that orthologues of the genes in non-model plants (e.g. maize) could simply be identified through homology searches and phylogenetic analysis. We first identified maize orthologues of *STR2* (putative lipid exporter) and *RAM2* that encodes glycerol-3-phosphate acyltransferase, a key enzyme of the AM-specific lipid biosynthetic pathway (32), and then analyzed their expression patterns, together with the mycorrhiza-inducible Pi transporter *Pht1;6* of maize (43, 44). Their expression levels were strongly correlated across the samples (SI Appendix, Fig. S1), which indicated that the expression of these genes are co-expressed and tightly regulated across ecosystems, irrespective of maize variety, and further, led us to the idea that plants have developed a gene regulatory unit specialized for mycorrhizal formation.

Based on this idea, we conducted weighted gene-coexpression network analysis on the transcriptome data of 251 samples using the R package with a soft-threshold power of 14 that provided a scale-free topology of the network (*r* ^2^ = 0.91) (SI Appendix, Fig. S2), resulted in definition of 19 coexpression modules. The 19 modules (named with colors) were functionally categorized according to the gene ontology (GO) enrichment analysis (Table 1 and Dataset S4): black, cell division; blue, transcription/translation; green, cell cycle regulation; greenyellow, stress-associated protein degradation; grey, immune response/N assimilation; grey60, antioxidation; lightcyan, branched-chain amino acid (BCAA) metabolism; lightgreen, water uptake and diurnal rhythm; magenta, immune response; midnightblue, lipid biosynthesis; pink, root development; royalblue, response to hypoxia; salmon, P-starvation response (PSR); tan, trehalose biosynthesis; yellow, mycorrhizal formation. No or only a few GO terms were enriched in cyan, darkgreen, darkred, and purple modules. The expression levels of these modules in individual samples are represented by eigengenes that are PC1 (sample) scores calculated based on the expression data of the module member genes (Dataset S5) (45). Pairwise correlation analysis of the eigengenes revealed that the grey, grey60, and lightcyan modules were positively correlated with each other (Fig. 2). In addition, these three modules were correlated positively with the same soil factors Bray II-P, NO_3_-N, Mg, and sand % (Dataset S6), suggesting that the three modules play general roles in nutrient assimilation/metabolism and immune responses. In contrast, the darkred module showed no distinct correlations with other modules nor with the soil/plant factors, indicating that the gene in the modules were expressed constitutively and unresponsive to environmental changes. Accordingly, this module is likely to have a “house-keeping” function. In subsequent analyses, we focused on the yellow (mycorrhizal formation), salmon (PSR), and pink (root development) modules that are likely to be involved directly in the nutrient acquisition strategies.

**Table 1.**
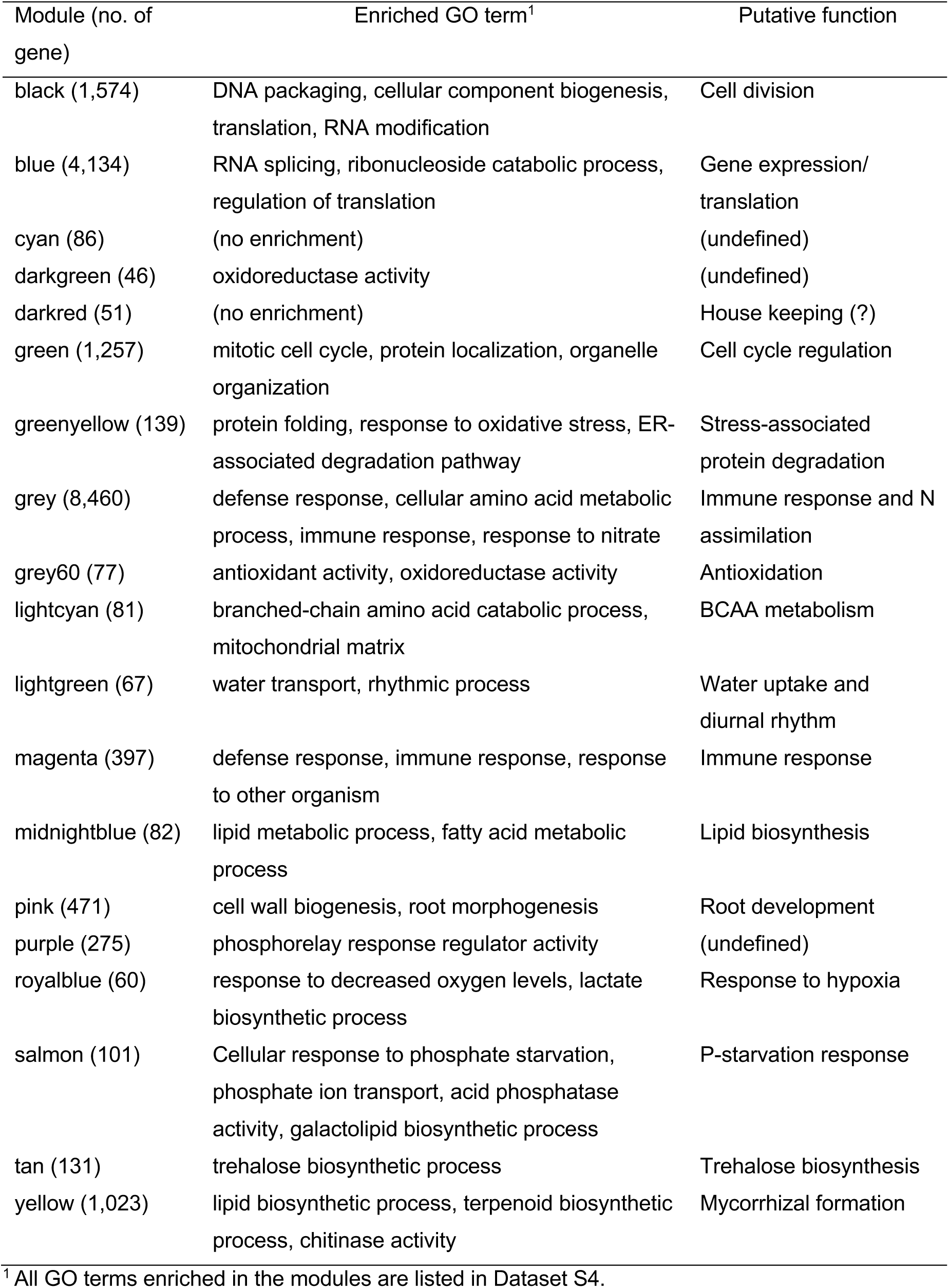
Enriched gene ontology (GO) terms and putative function of gene-coexpression modules.

| Module (no. of gene) | Enriched GO term<sup>1</sup> | Putative function |
| --- | --- | --- |
| black (1,574) | DNA packaging, cellular component biogenesis, translation, RNA modification | Cell division |
| blue (4,134) | RNA splicing, ribonucleoside catabolic process, regulation of translation | Gene expression/translation |
| cyan (86) | (no enrichment) | (undefined) |
| darkgreen (46) | oxidoreductase activity | (undefined) |
| darkred (51) | (no enrichment) | House keeping (?) |
| green (1,257) | mitotic cell cycle, protein localization, organelle organization | Cell cycle regulation |
| greenyellow (139) | protein folding, response to oxidative stress, ER-associated degradation pathway | Stress-associated protein degradation |
| grey (8,460) | defense response, cellular amino acid metabolic process, immune response, response to nitrate | Immune response and N assimilation |
| grey60 (77) | antioxidant activity, oxidoreductase activity | Antioxidation |
| lightcyan (81) | branched-chain amino acid catabolic process, mitochondrial matrix | BCAA metabolism |
| lightgreen (67) | water transport, rhythmic process | Water uptake and diurnal rhythm |
| magenta (397) | defense response, immune response, response to other organism | Immune response |
| midnightblue (82) | lipid metabolic process, fatty acid metabolic process | Lipid biosynthesis |
| pink (471) | cell wall biogenesis, root morphogenesis | Root development |
| purple (275) | phosphorelay response regulator activity | (undefined) |
| royalblue (60) | response to decreased oxygen levels, lactate biosynthetic process | Response to hypoxia |
| salmon (101) | Cellular response to phosphate starvation, phosphate ion transport, acid phosphatase activity, galactolipid biosynthetic process | P-starvation response |
| tan (131) | trehalose biosynthetic process | Trehalose biosynthesis |
| yellow (1,023) | lipid biosynthetic process, terpenoid biosynthetic process, chitinase activity | Mycorrhizal formation |

<sup>1</sup> All GO terms enriched in the modules are listed in Dataset S4.

**Figure 2.**
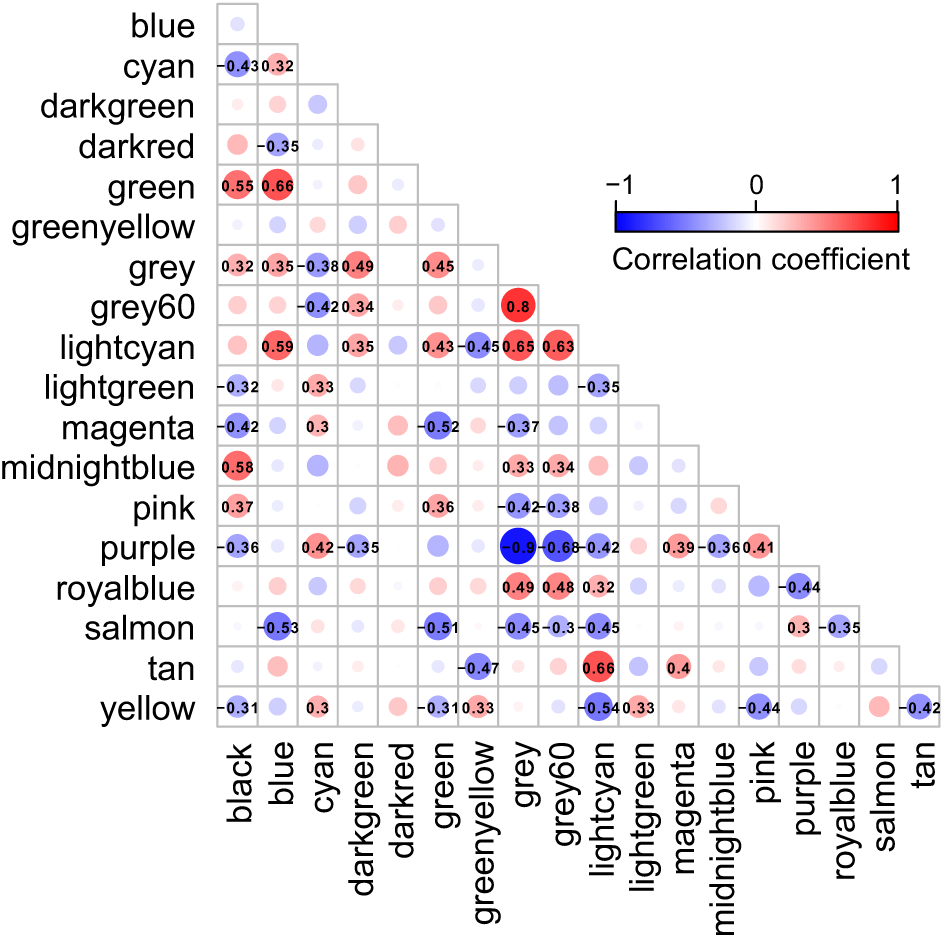
Correlation matrix of module eigengenes. Pairwise correlation analysis of the 19 coexpression modules defined by weighed gene coexpression network analysis was conducted, and correlation coefficients of which *P*-values are less than 0.001 are indicated. The blue and red circles indicate negative and positive correlations, respectively.

### Yellow module

A majority of the genes known to participate in arbuscule development and functioning are enriched in this module; e.g., the genes encoding the transcription factors RAM1 (46), RAD1 (47), and WRI5 (48), Pi transporter Pht1;6, nitrate transporter NPF4.5 (27), ammonium transporter AMT3;1 (28), H^+^-ATPase HA1 (49, 50), key enzymes for the mycorrhiza-specific lipid biosynthesis RAM2 and FatM (32), putative lipid exporters STR and STR2, and enzymes involved in strigolactone/mycorradicin biosynthesis DXS2 (51, 52), PSY2 (53), CCD7, CCD8, D27, and CCD1 (54) (Dataset S7).

To characterize this module in relation to AM fungal colonization, we analyzed unmapped sequence reads (i.e. the reads that were not mapped to the maize genome) that might contain AM fungal RNA reads. Generally, sequence reads of RNA-Seq (i.e. mRNA-Seq) contain a small number of those originated from ribosomal RNA (rRNA), which led us to the idea that AM fungal rRNA reads could be used as a fungal biomass index in the roots. We conducted Blastn searches against the large subunit rRNA sequences of maize and the AM fungal operational taxonomic units (OTUs) (55) using the unmapped reads as queries, and then fungal read counts for the OTUs were combined within each sample and normalized on the basis of plant rRNA read count (Dataset S8). It was considered, however, that fungal rRNAs that have A-rich regions were preferentially sequenced in mRNA-Seq because polyA-tailed RNA was purified prior to library construction, which might bias fungal rRNA abundance across the samples. To evaluate the significance of this bias, we sequenced rRNA directly (rRNA-Seq), which would provide unbiased rRNA read counts. Twenty RNA extracts were randomly chosen from the 251 samples and sequenced without purifying polyA-tailed RNA, and Blastn searches were performed using the rRNA reference sequences (Dataset S9). Correlation analysis showed that relative abundance of AM fungal rRNA read (normalized by plant rRNA reads) was highly correlated between the data obtained by the mRNA-Seq and rRNA-Seq (*r* = 0.919, *P* < 0.001) (SI Appendix, Fig. S3), suggesting that the bias is minimum. The unmapped reads were also *de novo* assembled, and the resulting 427,813 contigs (average length, 461 bp; N50, 525 bp) were subjected to Blastx searches against the 11 AM fungal (gromeromycotinan) genomes/transcripts (56-60) and 6 non-gromeromycotinan fungal genomes (SI Appendix, Table S3), as well as against the maize genome and Genbank bacterial genome database (SI Appendix, Fig. S4), to identify the AM fungal genes involved in Pi delivery, that is, those encoding putative Pi exporter SYG1-1, polyphosphate polymerase VTC4, vacuolar Pi exporter PHO91, and aquaglyceroporin AQP3 (24). We identified 86, 74, 159, and 105 contigs that showed similarity to the AM fungal *SYG1-1, VTC4, PHO91*, and *AQP3*, respectively, mapped the reads to these contigs, combined read counts within each gene, and normalized on the basis of TPM of the plant (Dataset S10). These expression data were subjected to multiple linear regression analysis using the yellow module eigengenes as an objective variable, together with the AM fungal rRNA read counts. Among the genes, the read counts of rRNA, *SYG1-1*, and *PHO91* were the significant explanatory variables for the eigengenes (SI Appendix, Table S2), suggesting that the eigengenes reflect not only fungal biomass but also the functionality of fungal colonization.

For further functional categorization of the 1,023 genes of this module, the k-means clustering analysis was performed using correlation distance as a measure. Clustering the genes into five submodules successfully separated into different functional groups along the PC2 axis of the yellow module (Fig. 3A and B and Table 2). In the submodule 1, a majority of the essential components of arbuscule development and nutrient exchange were enriched, e.g., *RAM1, RAD1, WRI5, Pht1;6, NPF4*.*5, AMT3;1, HA1, RAM2, FatM, STR*/*STR2*, serine/threonine receptor-like kinase ARK1 (61), and the genes involved in membrane trafficking *EXO70I* (62) and *Vapyrin A* (63) (Fig. 3C and Dataset S7). The enrichment of these genes suggests that the submodule 1 play a central role in the yellow module via modulating nutrient exchange across the periarbuscular membrane. In the submodule 2, the GO terms fatty acid biosynthetic process and plastid were overrepresented (Dataset S11), reflected in the enrichment of genes encoding the enzymes involved in the *de novo* biosynthesis of C16:0-fatty acid via acetyl-CoA in plastids. This suggests that the submodule 2 has a role in the biosynthesis of the essential component of the lipid for the construction of periarbuscular membrane and/or for the export to the fungi. In the submodule 3, the GO terms isoprenoid biosynthetic process, carotenoid metabolic process, and hormone metabolic process were overrepresented; the genes involved in the biosynthesis of carotenoid derivatives *DXS2, PSY2, CCD7, CCD8*, and *D27* were enriched. In addition, the GRAS transcription factor NSP2 that regulates strigolactone biosynthesis via modulating *D27* expression (64) also belongs to the submodule. Therefore, it is likely that the submodule 3 regulates the early processes of fungal accommodation through strigolactone production. In the submodule 4 the GO terms senescence-associated vacuole, amino sugar metabolic process, and extracellular space were overrepresented. The enrichment of the genes encoding a MYB1-like transcription factor, chitinase, and cysteine protease suggests that this submodule is responsible for arbuscule degeneration/turnover (65). Neither GO terms nor known genes involved in fungal accommodation/functioning were enriched in the submodule 5, and therefore, the biological function of this submodule could not be defined.

**Table 2.** Enriched gene ontology (GO) terms and putative function of the yellow submodules.

| Module (no. of gene) | Enriched GO term<sup>1</sup> | Putative function |
| --- | --- | --- |
| Submodule 1 (332) | peptidase activity, transferase activity | Nutrient exchange |
| Submodule 2 (203) | fatty acid biosynthetic process, plastid part | Fatty acid biosynthesis |
| Submodule 3 (219) | terpenoid biosynthetic process, carotenoid<br>metabolic process, plastid part | Carotenoid biosynthesis |
| Submodule 4 (189) | amino sugar metabolic process, defense<br>response to fungus, extracellular space | Arbuscule degeneration |
| Submodule 5 (80) | (no enrichment) | (undefined) |

<sup>1</sup> All GO terms enriched in the modules are listed in Dataset S11.

**Figure 3.**
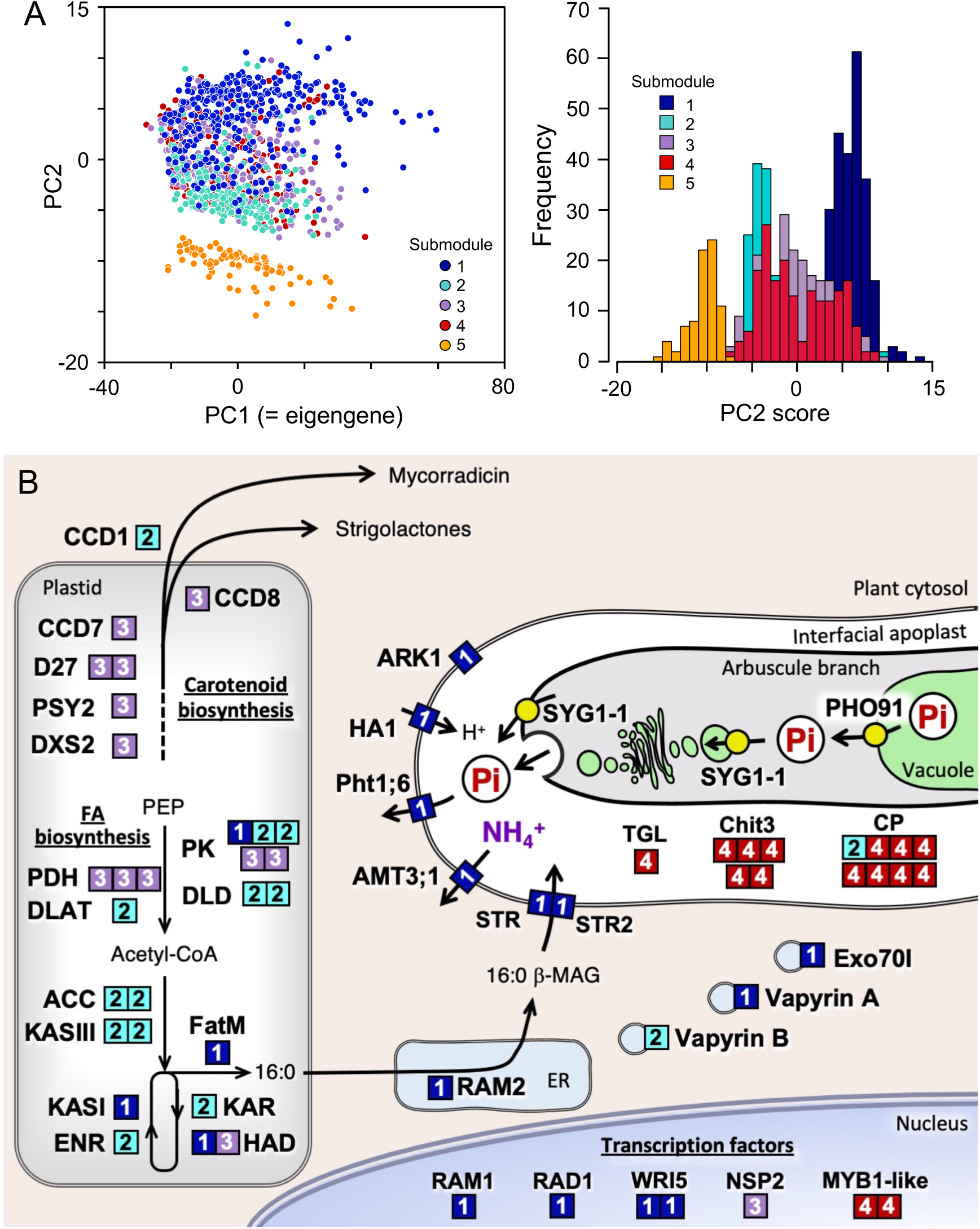
Functional categorization of the 1,023 genes of the yellow module into five submodules by k-means clustering analysis using correlation-based distance as a measure. (*A*) PCA plot of the genes in the submodules 1 – 5 (left), and frequency distribution of PC2 score of the genes (right). The submodule numbers the yellow module genes are indicated with the following colors: 1, dark blue; 2, turquoise; 3, purple; 4, red; 5, orange. (*B*) Putative function and cellular localization of the genes of which orthologues have been functionally characterized in previous studies. The numbers of submodule to which the genes belong are indicated in the boxes with the same colors in *A*. No orthologues of the genes in the submodule 5 has so far been functionally characterized, and thus these genes are excluded from this scheme.

Connectivities of the module member genes (i.e. correlation coefficients between their expression levels and the eigengenes) were calculated to assess the significance of the genes in the network (Datasets S7). In general, genes with a higher connectivity are defined as those that play a more significant role in the module, that is, “hub genes” (45). Average connectivities of the submodules 1 – 5 with the yellow eigengenes were 0.88, 0.72, 0.77, 0.75, and −0.59, respectively. In addition, most genes in the submodule 1 showed positive PC2 scores, while most genes in the submodules 2 and 5 showed negative PC2 scores (Fig. 3B), suggesting that these submodules are differentially regulated.

Taken together, the yellow (mycorrhizal) module consists of the five submodules that have different functions; the submodule 1 plays a central role in arbuscule development and nutrient exchange under the control of RAM1, RAD1, and WRI5, the submodule 2 is responsible for fatty acid biosynthesis for the construction of periarbuscular membrane and/or for carbon supply to the fungi, the submodule 3 is involved in carotenoid biosynthesis/metabolism, probably for the synthesis of strigolactones and mycorradicin, under the control of NSP2, and the submodule 4 is responsible for arbuscule degeneration/turnover under the control of MYB1-like transcription factor. Importantly, although the genes in this module are generally coregulated, the expression of submodules is finely tuned in response to environmental changes, which is analyzed in a later section.

### Salmon module

This module consists of 101 genes, in which the GO terms involved in PSR, e.g., cellular response to Pi starvation, Pi transport, acid phosphatase activity, and galactolipid biosynthetic process were overrepresented (Dataset S5). Pht1;3, one of the four Pi transporters in this module, is likely to be responsible for the direct pathway (66, 67) in which the transporter is localized to root epidermis and mediates Pi uptake independently of the mycorrhizal pathway (68). Six genes encoding SPX (SYG1-PHO1-XPR1) domain that is a sensor for the inositol polyphosphate-mediated signaling pathway for the maintenance of Pi homeostasis in eukaryotes (69) were also enriched in this module; one encodes the Pi exporter PHO1 (70) and the other five encode SPX-domain containing protein. Many acid phosphatase genes, including those encoding purple acid phosphatase (PAP), were assigned to this module. Two orthologues of *NIGT1* that encodes a MYB-type transcription factor belong to this module. NIGT1 was originally identified as a repressor of nitrate uptake (71), but also has a role in PSR via modulating the expression of SPX genes (72-74). It has been well documented that the transcription factor PHR1 plays a central role in PSR as a master regulator in Arabidopsis, a non-mycorrhizal plant (75). In maize two orthologues of PHR1 have so far been found (76), but both were assigned to the blue module in this study (Dataset S7), suggesting that, in the presence of mycorrhiza, PHR1 may not directly be involved in the regulation of PSR. In higher plants, including maize, Pi deficiency induces the replacement of membrane phospholipid with non-phosphorous lipids such as galactolipid and sulfolipid (77). The genes encoding glycerophosphodiester phosphodiesterase (GDPD2) for the degradation of phospholipid, monogalactosyldiacylglycerol (MGDG) synthase for the biosynthesis of galactolipid, and sulfoquinovosyl transferase (SQD2) for the biosynthesis of sulfolipid were assigned to this module. Higher connectivities with the eigengenes were observed for the genes encoding the enzymes involved in phospholipid replacement, MGDG synthase, SQD2, and GDPD2, SPX-domain containing protein, and NIGT1 (Datasets S7).

### Pink module

This module consists of 471 genes, and the GO terms involved in cell wall synthesis and root development, e.g., cell wall biogenesis, cellulose synthase, cytoskeleton, root system development, root morphogenesis, root epidermal cell differentiation, were overrepresented (Dataset S5), indicating that this module is responsible for the development of root system that accompanies transcriptional activation of a set of genes for cell wall biogenesis. Plant cell wall is composed of the primary cell wall that supports the fundamental growth of the cell and the secondary cell wall that supports the primary cell wall mechanically and facilitates water transport. CesA genes encode a cellulose synthase catalytic subunit and form cellulose synthase complexes consist of 18 – 24 *CesA* proteins; CesA1, CesA2, CesA3, CesA5, CesA6, and CesA9 are involved in the primary cell wall synthesis, while CesA4, CesA7, and CesA8 are for the secondary cell wall synthesis in Arabidopsis (78-80). In the maize genome twenty *CesA*s have been found (81), among which eleven genes, *CesA1, CesA2, CesA4, CesA6, CesA7a, CesA7b, CesA9, CesA10, CesA11, CesA12a*, and *CesA12b* are assigned to this module (Dataset S7). In addition to these *CesA*s, many genes encoding polysaccharide-biosynthetic/modification enzymes were found in this module; e.g., glycosyltransferases and glucanases. In this module there are 15 genes encoding the essential parts of cytoskeleton, alpha- and beta-tubulins and a microtubule motor protein. Eighteen transcription factors, e.g., an ethylene-responsive transcription factor (ERF), five NAC-domain transcription factors, and seven MYB-domain transcription factors, were also found. ERF035 was expressed in regions related to cell division/differentiation, e.g., lateral roots and central cylinder of primary roots, and upregulate *CesA1* in Arabidopsis (82). It has been known that many NAC- and MYB-domain transcription factors are responsible for the regulation of cell wall synthesis and expressed during root formation (83-86). Connectivities between the eigengenes were higher in those encoding cellulose synthases, polysaccharide modification enzymes, and microtubule-related proteins (Dataset S7).

### Module interplays

For understanding interplays between the modules in relation to the soil and plant factors, multiple regression analysis was conducted, in addition to simple linear regression analysis. Basically, the eigengenes of yellow and salmon modules were not correlated (Figs. 2 and 4A). However, the expression levels of 65 out of the 101 salmon genes were positively correlated with the yellow eigengenes (*P* < 0.05), which include three out of the four Pi transporter genes and all eight acid phosphatase genes (Fig. 4B and Dataset S7). Furthermore, most genes in the submodules 1 (arbuscule development and nutrient exchange) and 3 (carotenoid biosynthesis and metabolism) of the yellow module were also positively correlated with the salmon eigengenes (*P* < 0.05). The yellow eigengenes were mainly explained by the plant factors leaf P content (with a negative coefficient), leaf P:N ratio (positive), stem diameter (positive), and growth rate (negative) and the soil factors NO_3_-N (negative), clay % (negative), and Bray II-P (negative) in the multiple regression analysis (Fig. 4C). In addition to these factors, leaf N content was also negatively correlated with the eigengenes in the simple linear regression analysis (Dataset S6). The salmon eigengenes were largely explained by the plant factor growth rate (negative) and leaf P:N ratio (negative) and the soil factors organic matter % (positive), Bray II-P (negative), exchangeable K (positive) and Ca (negative), and silt % (positive) both in the multiple regression and simple linear regression analyses. These results suggest that, although the two modules are regulated independently, parts of the genes in these modules are coordinately expressed, particularly under P-deficient conditions. It is likely, however, that N deficiency is also an important driver for the yellow module, given that N deficiency leads to increases in leaf P:N ratio and decreases in leaf N content that are positive and negative factors, respectively, for the module.

**Figure 4.**
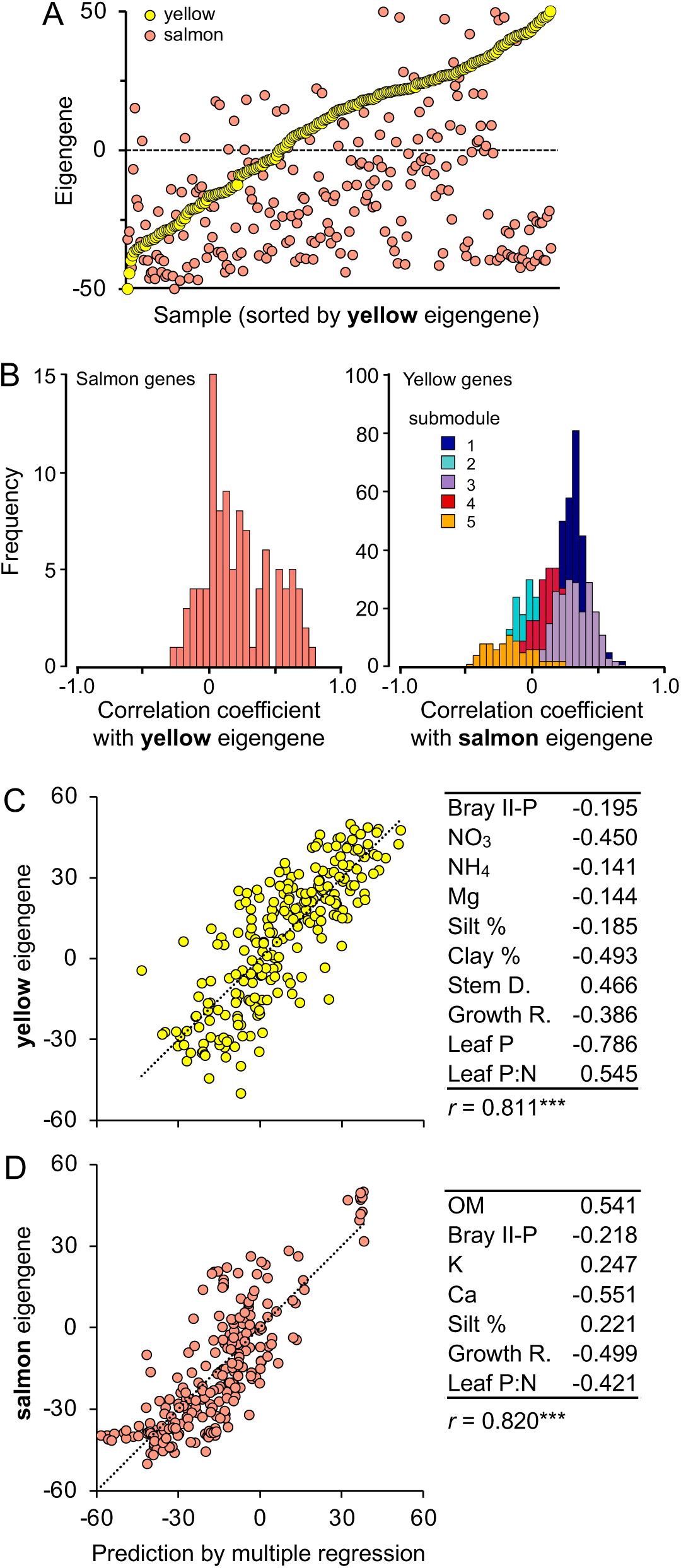
Interplay between the yellow and salmon modules in relation to soil and plant factors. (*A*) Scatter plot of yellow (yellow) and salmon (salmon) eigengenes of the 251 samples, in which the samples were sorted by the order of yellow eigengene. The eigengenes were standardized between the module from −50 (minimum value) to +50 (maximum value). (*B*) Frequency distributions of correlation coefficients of the salmon module genes with the yellow eigengenes (left) and those of the yellow module genes with the salmon eigengenes (right). The submodule numbers of the yellow module genes are indicated with the following colors: 1, dark blue; 2, turquoise; 3, purple; 4, red; 5, orange. The data were extracted from Dataset S6. Multiple regression analysis of the yellow (*C*) and salmon (*D*) eigengenes with the soil and plant factors. Regression coefficients of the factors with *P* < 0.001 are indicated in the tables. The correlation coefficients are also indicated below the tables (***, *P* < 0.001).

Plant availability of Pi in soil is generally too low to meet the plant demand because most P is present as unavailable forms, such as sparingly soluble Pi and organic P (87). Exploitation of the unavailable P by the secretion of organic acids and phosphatase, that is, typical PSRs, and mycorrhizal formation are two major strategies for the acquisition of Pi, but their interplays have been largely overlooked. Although the salmon (PSR) and yellow (mycorrhizal) modules similarly respond to many plant and soil factors, their responses to leaf P:N ratio is contrasting; the factor is a negative regulator for the PSR module but a positive regulator for the mycorrhizal module. The upregulation of PSR by lower leaf P:N ratio (i.e. P deficiency relative to N) could be interpreted by the high-connectivity of *NIGT1* with the salmon eigengenes because *NIGT1* that is involved in the signaling cascade of PSR (71, 74) is upregulated in a nitrate concentration-dependent manner (71). Furthermore, the fact that this regulatory cascade is conserved not only in the non-mycorrhizal plant Arabidopsis but also in the mycorrhizal plant maize strongly suggests that the PSR module has evolved independently from the mycorrhizal module. This independency of the PSR module would facilitate an alternative strategy to cope with Pi deficiency when Pi delivery through the mycorrhizal pathway does not meet the P demand due to, e.g., low-population density or absence of AM fungi and extremely low-Pi availability in the soil. On the other hand, the coexpression of the submodules 1 and 3 and some of the PSR genes suggests the presence of a common regulatory pathway for these genes that has to be elucidated.

Intriguingly, the eigengenes of yellow and pink modules were negatively correlated (Figs. 2 and 5A), which is reflected in negative correlations of the expression levels of the genes of the pink module with the yellow eigengenes and those of the genes of the yellow submodules 1-4 with the pink eigengenes (Fig. 5B). The pink eigengenes were explained by the plant factors leaf P content (with a positive coefficient) and leaf P:N ratio (negative) and the soil factor clay % (positive) in the multiple regression analysis (Fig. 5C). In addition to these factors, leaf N content were positive factors for the expression in the simple linear regression analysis (Dataset S6). These results indicate that the pink module responds to the plant and soil factors in the opposite direction to which the yellow module responds, suggesting that root development is intrinsically an opposite strategy of mycorrhizal formation.

**Figure 5.**
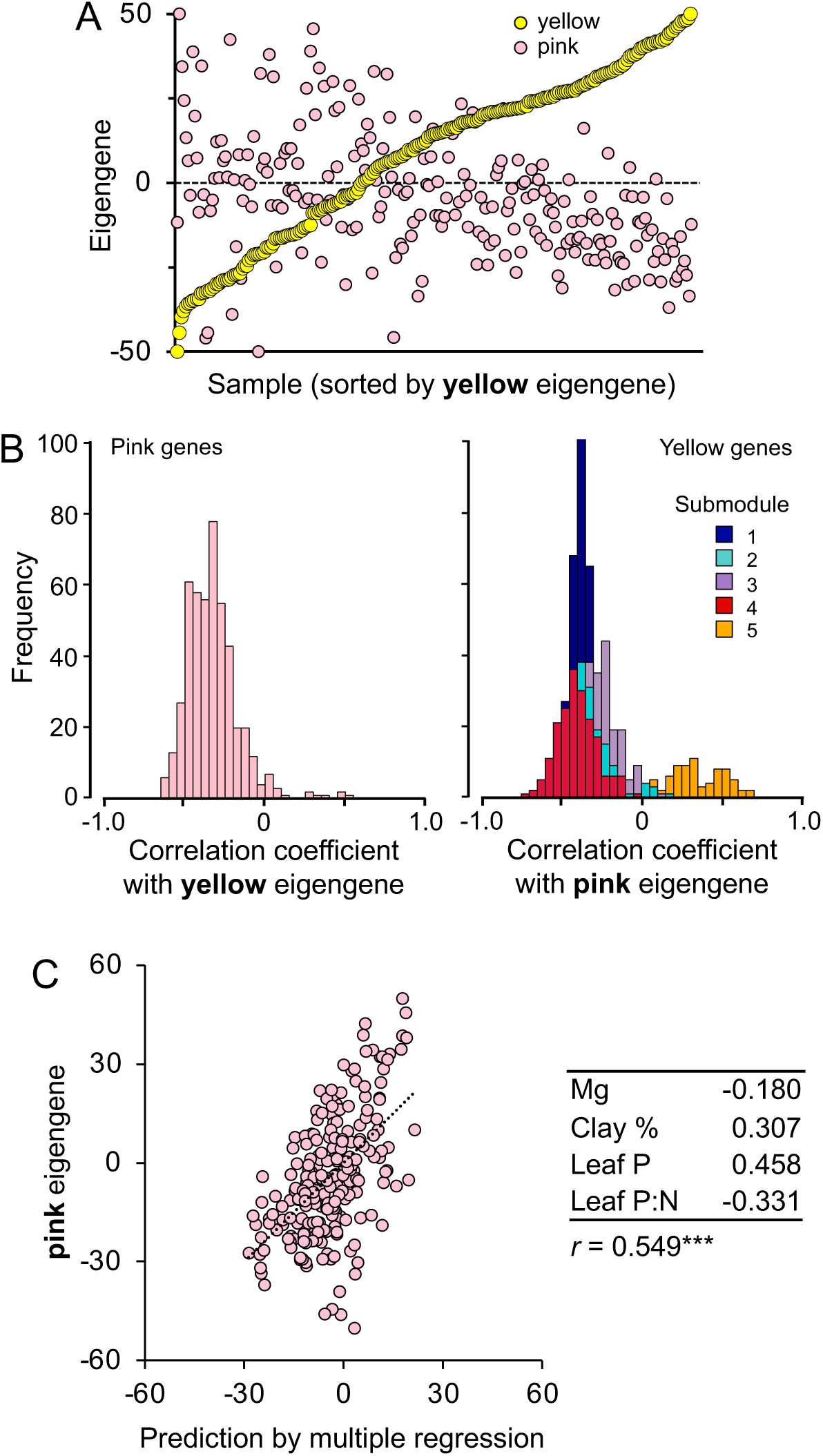
Interplay between the yellow and pink modules in relation to soil and plant factors. (*A*) Scatter plot of yellow (yellow) and pink (pink) eigengenes of the 251 samples, in which the samples were sorted by the order of yellow eigengene. The eigengenes were standardized between the module from −50 (minimum value) to +50 (maximum value). (*B*) Frequency distributions of correlation coefficients of the pink module genes with the yellow eigengenes (left) and those of the yellow module genes with the pink eigengenes (right). The submodule numbers of the yellow module genes are indicated with the following colors: 1, dark blue; 2, turquoise; 3, purple; 4, red; 5, orange. The data were extracted from Dataset S6. (C) Multiple regression analysis of the pink eigengenes with the soil and plant factors. Regression coefficients of the factors with *P* < 0.001 are indicated in the table. The correlation coefficient is indicated below the table (***, *P* < 0.001).

In the Early Devonian, AM fungal symbiosis facilitated terrestrialization of early plants that did not have functional root systems by providing a water/nutrient uptake pathway (88), and during the Middle to Late Devonian, plants evolved a substantial root system that is capable of not only taking up water/nutrients but also anchoring the body to the soil (89). In the context of water/nutrient uptake, mycorrhizal formation and root development are functionally redundant. We demonstrated that the two strategies are oppositely regulated as a function of plant nutrient status, that is, plants allocate more resource to mycorrhizal formation under nutrient-deficient conditions, but under nutrient-enriched conditions, more resource to root development. This could be interpreted by a trade-off between cost for the extension of surface area for nutrient uptake and the rates (efficiency) of nutrient uptake and translocation. The mycorrhizal pathway is mediated by fungal hyphae that are finer than roots, providing a larger surface area per carbon investment than root system development. It is likely, therefore, that the mycorrhizal pathway is more efficient under nutrient-depleted conditions in which nutrient uptake and translocation by hyphae are more rapid than the diffusion rates of nutrients towards the roots (68). On the other hand, the root direct pathway may facilitate more rapid nutrient uptake and translocation under nutrient-enriched conditions in which the diffusion rates are rapid enough to sustain the uptake from the pathway. In addition, root development would play an important role in physical support for the above-ground part that grows vigorously under nutrient-enriched conditions. It has been suggested that gramineous plants have evolved finer roots to obtain a larger surface area per unit carbon investment, that is, towards less dependency to mycorrhizas (90). Our findings suggest, however, that plants have also developed a regulatory mechanism for modulating resource allocation between mycorrhizas and a root system to optimize the efficiency of nutrient acquisition in a given environment.

## Conclusion

Our study provides a new insight into the regulatory mechanism of plant nutrient-acquisition strategies in complex environments. We revealed the distinct genetic modules for nutrient acquisition strategies in maize; that is, mycorrhizal formation, PSR, and root development and, further, identified soil and plant factors that drive these modules. The expression levels of the mycorrhizal module were correlated not only with fungal biomass (i.e. rRNA abundance) but also with the expression levels of fungal genes involved in nutrient delivery, reflecting the functionality of mycorrhiza. The two distinct responses to P deficiency, the enhancement of P acquisition via the root direct pathway and the internal recycling of P, are coregulated at the transcriptional level in the PSR module. The root development module that consists of a large number of the genes involved in cell wall biogenesis is upregulated under nutrient-enriched conditions, suggesting that this strategy requires substantial carbon investment (e.g., 91).

In maize production, different genotypes are grown in different regions, and a number of new genotypes are distributed every year. In this study, a variety of genotype was collected across the countries and analyzed without taking into account genotypic differences. It has been well documented that different genotypes respond to biotic and abiotic factors differently; for instance, Pi acquisition efficiency via the mycorrhizal pathway is different among maize genotypes (43, 92). We consider, however, that the genetic modules identified in this study are conserved across genotypes of the species and that such differences in the responses are due to differential regulation of the expression of the modules among the genotypes. This issue could not be addressed in our current approach and has to be elucidated in future studies.

Our cross-ecosystem transcriptomics approach opens a new window for understanding how plants respond to complex environments, providing implications for sustainable management of agriculture. We identified the soil factors that drive the modules for nutrient acquisition, which suggests that resource allocation to the nutrient uptake pathways and PSRs could be optimized via manipulating these factors, e.g., via balancing fertilizer input and improving soil physical conditions. Towards sustainable agriculture, replacement of chemical fertilizer (i.e. non-renewable resource) with renewable resource, e.g., organic matter, is also necessary. In this study, we focused only on conventional agriculture in which soil fertility was managed mainly by chemical fertilizer. Further studies are needed to clarify how to balance resource allocation to the strategies in the crop in organic agriculture.

## Materials and Methods

The plant and soil samples were collected from 21-to 54-day-old plants from the USA in June 2017 and from 13-to 54-day-old plants from Japan in May – July 2018. Details of geographic, climatic, and field-management data of the sampling sites/plots are presented in SI Appendix, Table S1. Field sampling, soil and plant analysis, RNA sequencing, gene expression profiling, rRNA read assignment, *de novo* assembling of AM fungal mRNA reads, and statistics were performed as described in SI Appendix, Supplementary Information Text.

## Data availability

The RNA-Seq reads have been deposited in National Center for Biotechnology Information under the accession number PRJNA586746 (mRNA-Seq, 251 samples) and PRJNA604657 (rRNA-seq, 20 samples). The expression data of the maize roots and AM fungal transcript references can be downloaded from: http://lab.agr.hokudai.ac.jp/botagr/rhizo/RhizoCont/Download.html.

## Acknowledgments

The authors acknowledge A. Nagano for technical advice, S. Inman and S. Garcia for the arrangement of field experiments and sampling in the USA, T. Mitani, K. Tawaraya, T. Koyama, Y. Tahara, and T. Nagaoka for the arrangement of field experiments and sampling in Hokkaido University, Yamagata University, Utsunomiya University, Nagoya University, and Hiroshima University, respectively. This work was partially supported by ACCEL (JPMJAC1403) form Japan Science and Technology Agency (Y.S., H.M, and TE).

## Supplementary Information for

### Other supplementary materials for this manuscript include the following

Datasets S1 to S11

## Supplementary Information Text

### Field sampling and soil/plant analysis

The root system was collected from a 30 × 30 cm area at a depth of 20 cm and washed briefly and gently in water. Then primary, lateral, and seminal roots (except for crown roots) were immediately collected with scissors, cut into small pieces (< 1 cm), randomized in water, collected on a stainless mesh, and blotted with a paper towel, and about 10 mg of the root pieces were immersed in 15 mL of RNAlater solution (Thermo Fisher Scientific), which were done within 3 min for each sample in the field sites. Fully-expanded-youngest leaves were also collected from the same individuals for P and N analysis. Leaf number, stem diameter (1 – 2 cm-above ground), and dry weight of the above ground part of the same individuals were measured. Root-zone soil samples (approx. 500 g) were also collected form the same individuals for the analysis of physical and chemical properties. The root samples were kept in RNAlater at least for 48 h at room temperature, transferred onto a stainless mesh to remove excess RNAlater, blotted, and stored in plastic tubes below −80°C.

Chemical and physical properties of the US and Japanese soil samples were analyzed at Midwest Laboratories (Omaha, NE) and Tokachi Federation of Agricultural Cooperatives (Obihiro, Japan), respectively, by the standard methods. The maize above-ground part and leaf samples were dried at 80°C for up to 5 days and weighed. The leaf samples were ground and digested with concentrated sulfuric acid and hydrogen peroxide at 200°C for 150 min, and then N and P concentrations were determined by the indophenol-blue method (1) and ascorbic acid-molybdate blue method (2), respectively.

### RNA sequencing and gene expression profiling

Total RNA was extracted from the root samples using Maxwell RSC Plant RNA Kit with Maxwell RSC Instrument (Promega). Sequencing libraries were constructed using KAPA Stranded mRNA-Seq Kit (KAPA Biosystems), and single-end 75-base sequencing was performed on the illumina NextSeq 500 platform (10 M reads per sample) at Bioengineering Lab, Sagamihara, Japan. In ribosomal RNA sequencing (rRNA-Seq), the sequence libraries were constructed without mRNA purification. The sequence reads of which Phred quality score ≥20 in >80% of the bases were mapped to the maize B73 reference genome sequence (Zm-B73-REFERENCE-GRAMENE-4.0, ftp://ftp.ensemblgenomes.org/pub/plants/release-36/fasta/zea_mays/dna/) using HISAT2 program (3). Mapped reads were extracted and converted into BAM format using SAMtools (4), and uniquely mapped reads to the protein-coding transcript sequences were counted with featureCounts program (5). The raw read counts were normalized to Transcripts Per kilobase Million (TPM), and then genes with an average TPM ≥ 5 were subjected to subsequent analyses.

To identify maize genes involved in mycorrhizal formation and functioning, Blastp searches were performed against the maize genome using the amino acid sequences of those identified in *Oryza sativa, Sorghum bicolor, Medicago truncatula*, and *Lotus japonicus* as queries, and candidate genes were subjected to phylogenetic analysis based on the Neighbor-Joining method with MEGA X (6).

For weighted gene coexpression network analysis (WGCNA), the TPM values of the genes were log_2_-transformed and analyzed using the R package WGCNA with the settings of ‘automatic, one-step network construction’, mergeCutHeight of 0.2, and minimal module size of 30 genes. After definition of modules, we merged the modules with a distance threshold value of 0.3. The k-means clustering analysis was performed using correlation distance as a measure with the R package amap. Fisher’s exact test was employed to analyze GO term enrichment using Blast2GO program with false discovery rates of less than 0.01 and 0.05 for the modules and submodules, respectively (7). The GO annotation list was obtained from the maize genome database.

### rRNA read assignment and *de novo* assembling of AM fungal mRNA reads

Maize and AM fungal rRNA sequences in the rRNA-Seq and the corresponding mRNA-Seq data were assigned to the maize large-subunit (LSU) rRNA sequence (XR_002749536) and 524 operational taxonomic units (OTUs) of AM fungal LSU rRNA sequences by Blastn searches with the following parameters: similarity, ≥ 95%; minimum alignment length, ≥ 75 bp; E-value, ≤ 1e-30. The consensus sequences GTGAAATTGTTGAAAGGGAAACG and GACGTAATGGCTTTAAACGAC at the 5’ and 3’ ends, respectively, of the AM fungal sequences were removed before analysis. The numbers of the reads assigned to the plant rRNA and AM fungal OTUs were normalized to unit nt length, and total AM fungal read counts in the samples were standardized per 10^5^ plant rRNA reads and transformed to logarithmic values.

For *de novo* assembling of AM fungal mRNA reads, the sequencing reads that were not mapped to the maize genome were collected from all 251 samples, combined into a fastq file, and a half of the reads was randomly extracted twice using SeqKit toolkit (8). The resultant two read sets were assembled separately with Trinity (ver. 2.4.0) (9), and then the contigs generated from the two batches were combined and clustered using CD-HIT-EST (10, 11) with a 100%-identity cutoff, and subjected to Blastx searches at an E-value cutoff of 1e-5 against the genomes/transcripts of *Rhizophagus irregularis* DAOM181602, *R. clarus* HR1 (MAFF520076), *Diversispora epigaea* IT104, *Gigaspora margarita* BEG34, *Funneliformis mosseae* DAOM236685, *Acaulospora morrowiae* INVAM-CR315B, *Diversispora versiforme* INVAM-W47540, *Scutellospora calospora* INVAM-IL209, *Racocetra castanea* BEG1, *Paraglomus brasilianum* DAOM240472, *Ambispora leptoticha* INVAM-JA116, *Saccharomyces cerevisiae* S288c, *Neurospora crassa* OR74A, *Laccaria bicolor* S238N-H82, *Ustilago maydis* 521, *Rhizopus oryzae* 99-892, and *Phycomyces blakesleeanus* NRRL1555 (Table S3), as well as against the maize genome and Genbank bacterial genome database, in which contigs that showed the highest similarity to AM fungal sequences were considered as those originated from AM fungi. From the AM fungal contigs, those containing an open reading frame (ORF) of 50 or longer amino acid residues were predicted using TransDecoder (https://transdecoder.github.io). These processes are summarized in Fig. S4. The high-quality mRNA reads were mapped to the ORF sequences as described in the previous section.

### Statistics

Principal component analysis and multiple linear regression analysis were performed using the prcomp and lm functions, respectively, in R. In variable selection in these analyses, one of the variables (soil and plant factors) that were highly correlated (|*r*| ≥ 0.9) were excluded (Dataset S2). In addition, multicollinearity was also considered in multiple regression analysis. Prior to the multiple regression analysis between eigengenes and soil/plant factors, all variables were standardized between −50 (minimum) and +50 (maximum).

**Fig S1.**
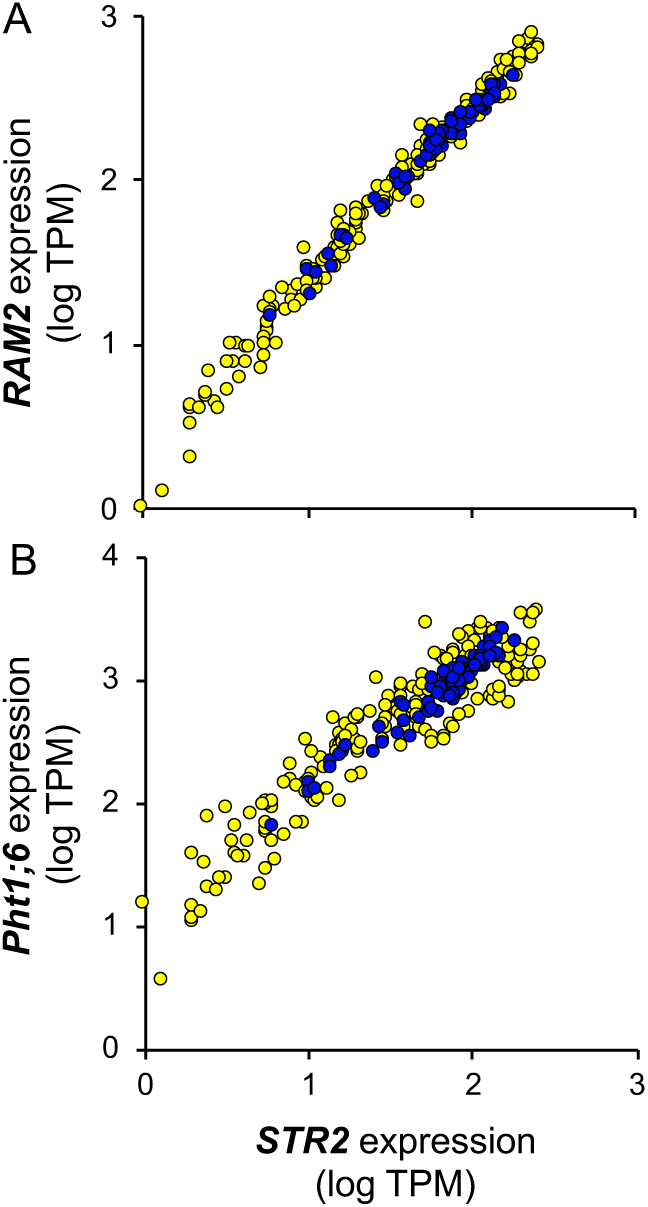
Scatter plots of the gene expression levels of the Pi transporter (*Pht1;6*), putative lipid transporter (*STR2*), and glycerol-3-phophoshate acyltransferase (*RAM2*) that are essential for arbuscule development and functioning. Correlation coefficients between *RAM2* and *STR2* and between *Pht1;6* and *STR2* are 0.939 and 0.930, respectively (*P* < 0.001).

**Fig S2.**
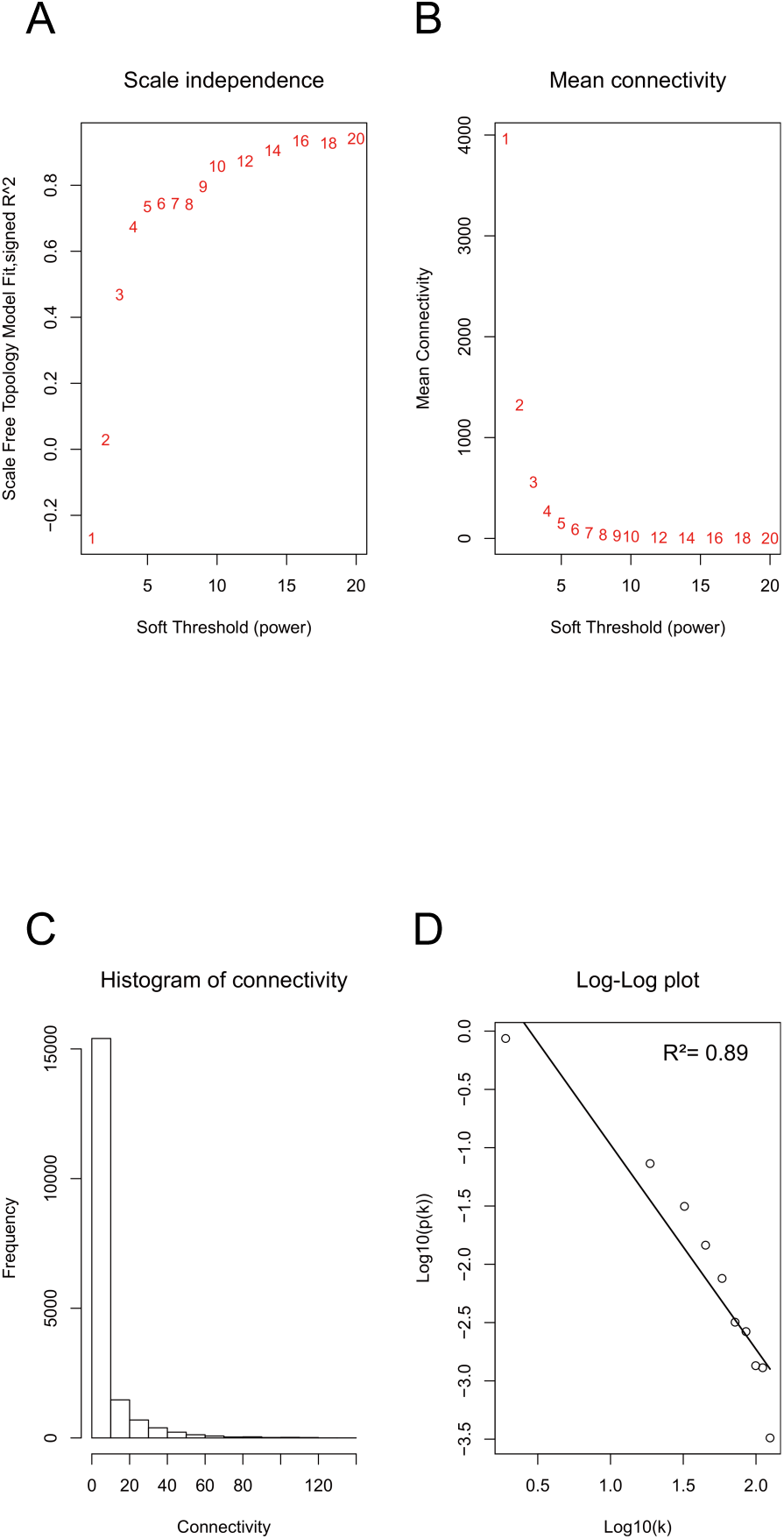
Evaluation of weighted gene coexpression network properties in maize roots. (*A*) Scale-free fit index as a function of soft-threshold power. (*B*) Mean connectivity as a function of soft-threshold power. (*C*) Frequency distribution of network connectivities. (*D*) Log-log plot of edges [log10(k)] versus probability of a node having k edges [P(k)].

**Fig S3.**
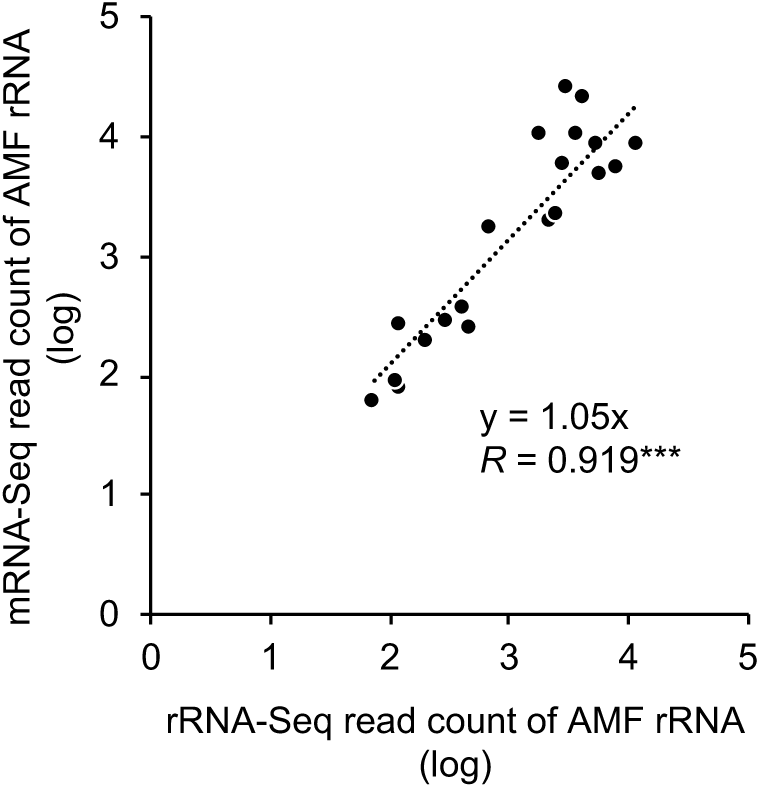
Correlation analysis between AM fungal rRNA read numbers in the mRNA-Seq and those in the rRNA-Seq. Twenty samples were randomly chosen from the 251 RNA samples and subjected to rRNA-Seq (i.e. RNA-Seq without purification of mRNA). The reads obtained by mRNA-Seq and rRNA-Seq were assigned to maize LSU rRNA and 524 AM fungal operational taxonomic units by Blastn searches. The read counts were normalized to unit nt length, and total AM fungal read counts in the individual samples were standardized by 10^5^ plant rRNA reads and transformed to logarithmic values.

**Fig S4.**
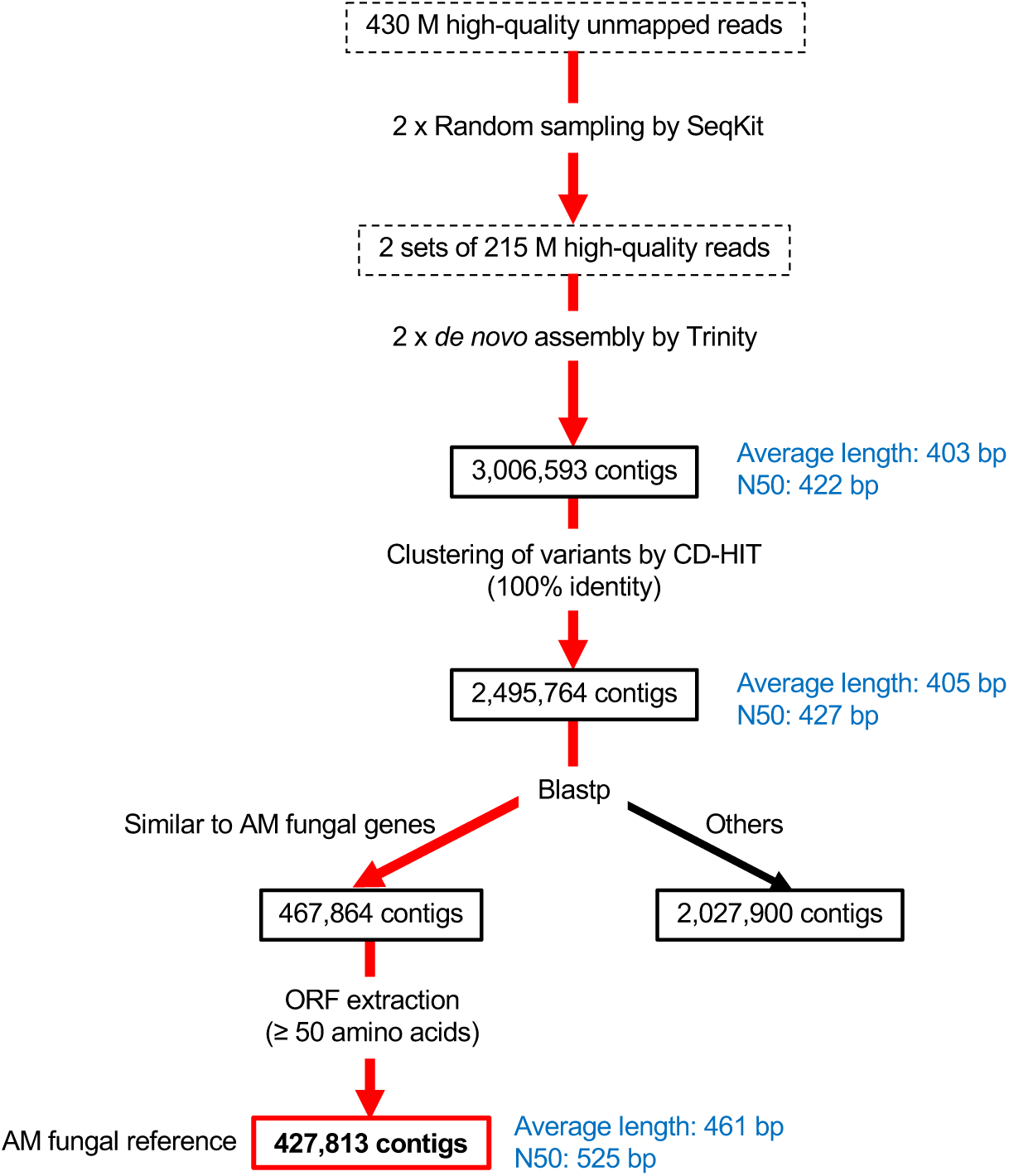
Procedure for the construction of AM fungal transcript references from the unmapped reads of 251 samples.

**Table S1.**
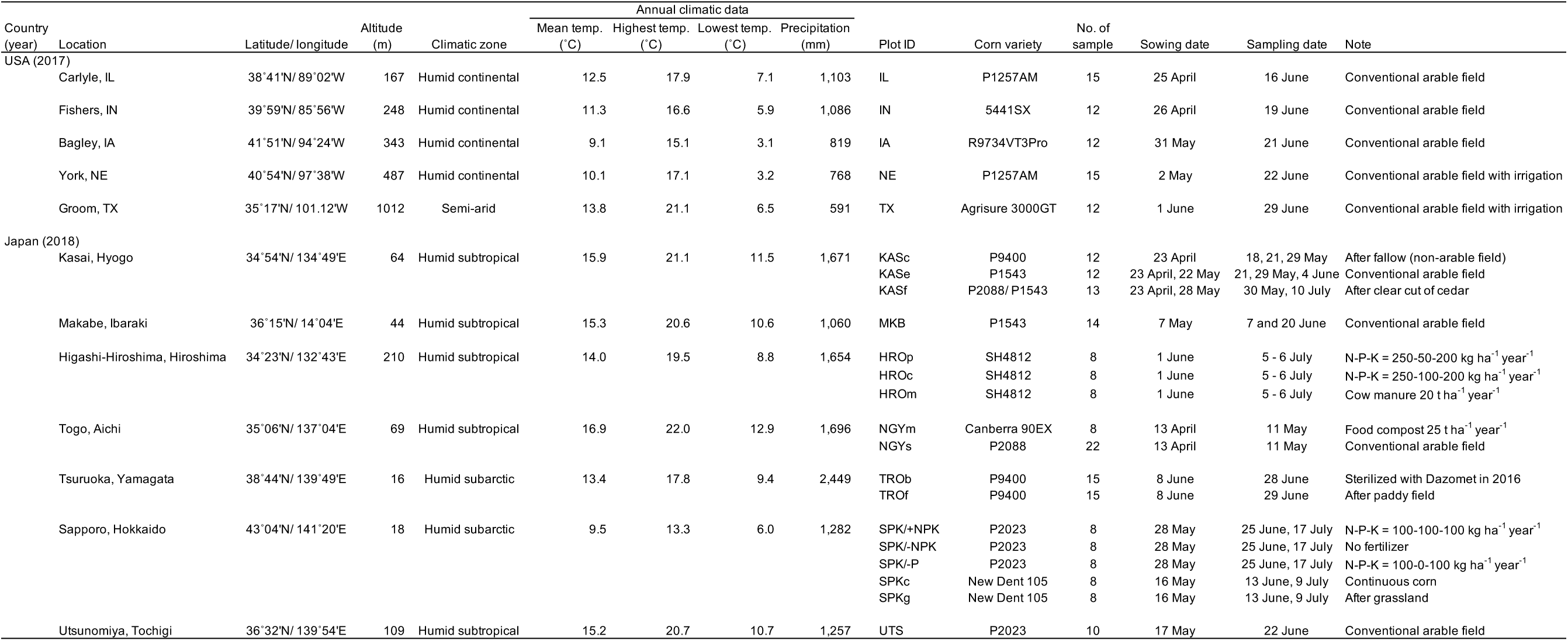
Geographic, climatic, and management data of the sampling sites.

**Table S2.** Multiple regression analysis of the yellow module eigengenes with the transcript levels of AM fungal genes.

|  | Regression<br>coefficient |  |
| --- | --- | --- |
| *SYG1-1* expression | 0.150 | *** |
| *PHO91* expression | 0.233 | ** |
| rRNA abundance | 0.875 | *** |
| Correlation coefficient (*r*) | 0.884 | *** |

\*\*, $P < 0.01$ ; \*\*\*, $P < 0.001$

**Table S3.**
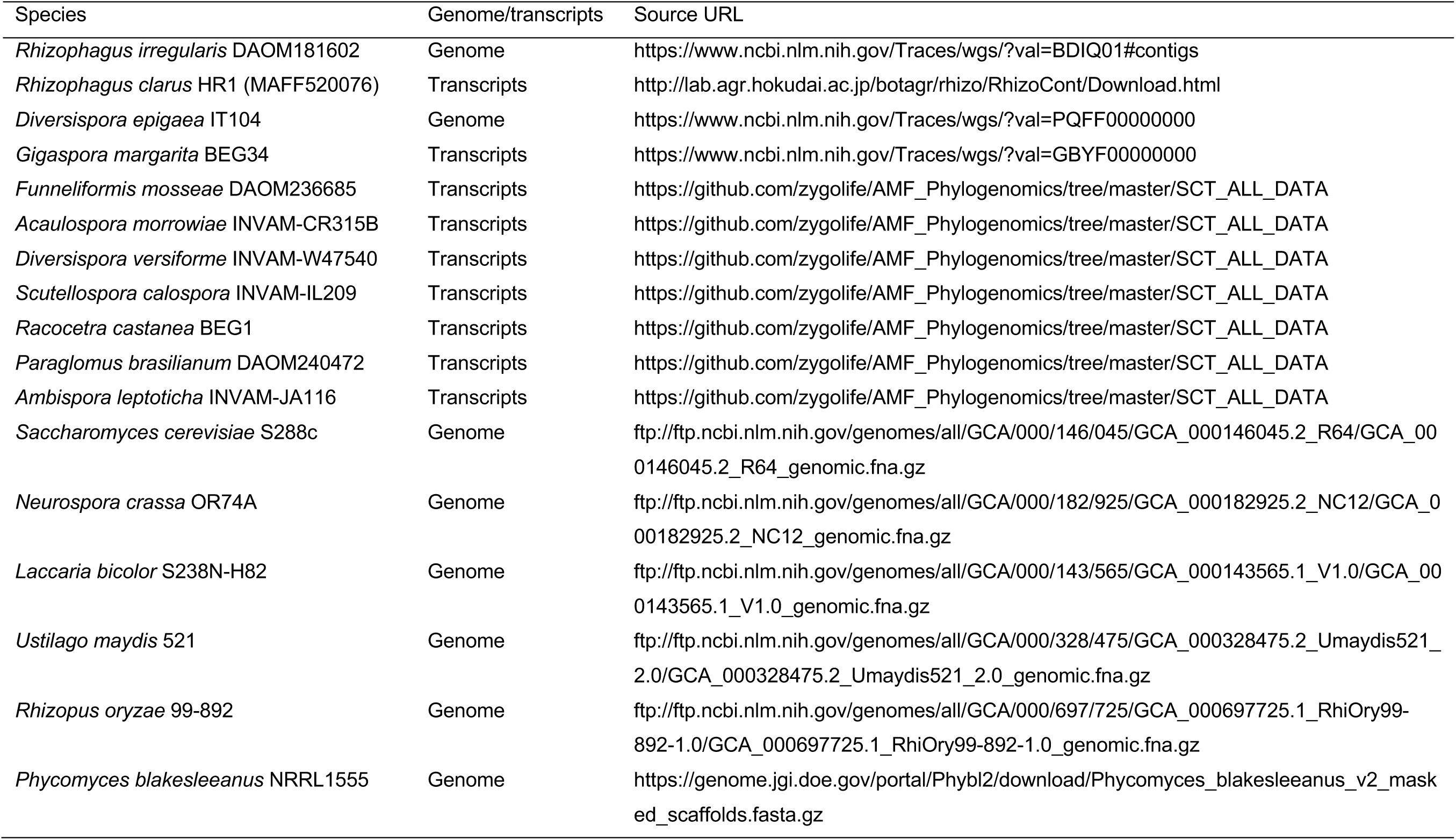
Source of fungal databases for Blastx searches of the contigs obtained by the *de novo* assembly of the unmapped reads.

**Dataset S1**. Soil and plant factors used for principle component analysis and correlation analysis.

**Dataset S2**. Correlation coefficient and probability between soil and plant factors.

**Dataset S3**. Read assignment summary.

**Dataset S4**. Gene ontology (GO) analysis of the coexpression modules in the maize roots.

**Dataset S5**. Eigengenes of the coexpression modules in the maize roots.

**Dataset S6**. Regression coefficients in correlation analysis between the eigengenes and soil/plant factors.

**Dataset S7**. Correlation coefficients and probabilities between the expression levels of individual genes and the eigengenes.

**Dataset S8**. Number of mRNA-Seq read assigned to the LSU rRNA genes of AM fungal operational taxonomic units and maize.

**Dataset S9**. Number of rRNA-Seq read assigned to the LSU rRNA genes of AM fungal operational taxonomic units and maize in randomly selected 20 samples.

**Dataset S10**. Expression levels (TPM) of AM fungal *SYG1-1, PHO91, VTC4*, and *AQP3* of which transcript references were obtained by *de novo* assembling of unmapped reads.

**Dataset S11**. Gene ontology (GO) analysis of the submodules of yellow module.

